# Y-linked editors for invasive rodent control

**DOI:** 10.64898/2025.12.08.693037

**Authors:** Prateek Verma, Omar S. Akbari, John M. Marshall

**Affiliations:** Divisions of Biostatistics & Epidemiology, School of Public Health, University of California, Berkeley, California, United States of America; School of Biological Sciences, Department of Cell and Developmental Biology, University of California, San Diego, California, United States of America; Innovative Genomics Institute, Berkeley, California, United States of America

**Keywords:** Biosafety, gene drive, genetic biocontrol, *Mus musculus*, native species conservation, population dynamics, population suppression, self-limiting, sterile male release

## Abstract

Invasive rodents are major contributors to biodiversity loss, particularly on islands where native species have evolved in their absence, rendering them vulnerable to predation. Genetic biocontrol offers a promising suite of species-specific technologies that may contribute to rodent suppression and elimination. Here, we evaluate the potential of Y-linked genome editors (YLEs), which contain a nuclease that disrupts a female-essential fertility or viability gene, to suppress or eliminate a mouse population on a small, isolated island. We benchmark YLEs against other self-limiting genetic biocontrol tools, such as sterile male releases, and self-sustaining tools, such as population suppression gene drives and Y-linked X chromosome-shredders. We use the MouseGD simulation framework to model the inheritance patterns of these systems in the context of rodent life history, calculating elimination probabilities, times to elimination and other outcomes for a range of construct designs, fitness costs and release schemes. We find that YLEs are more efficient than other self-limiting tools, and are capable of achieving rodent elimination within a short timeframe for modest release sizes. For a mouse population size of 10,000, elimination is predicted within five years for releases of 350 males per month, and within ten years for releases of 150 males per month. This scale of production is well within existing capabilities, potentially enabling suppression to encompass a larger spatial scale. We found that elimination could be achieved for YLEs targeting both haploinsufficient and haplosufficient target genes; but for a more restrictive parameter space for the haplosufficient case. Gene drives were predicted to achieve suppression and elimination for smaller release sizes; but also to spread to non-target populations. In contrast, YLEs do not bias inheritance and hence display minimal spillover. Altogether, these characteristics present YLEs as a promising, ecologically-manageable biocontrol tool for elimination of invasive rodents and conservation of native species.

## 1. Introduction

Islands are global biodiversity hotspots, supporting rich assemblages of species that have evolved in long-term isolation, many of which are endemic to these environments [1,2]. Introduction of alien invasive species poses profound threats to these vulnerable ecosystems. Among them, rodents (rats, in particular) are especially destructive; being responsible for ~40-60% of bird and reptile extinctions on islands [3,4]. In Aotearoa New Zealand, invasive predators, including stoats, rats and possums, kill approximately 25 million native birds each year, and roughly 4,000 native species are currently threatened or at risk of extinction [5]. Beyond ecological impacts, invasive rodents impose substantial economic costs, with conservative global estimates reaching US $87.5 million annually between 1980 and 2022 [6]. Effective rodent eradications can yield substantial and immediate conservation benefits [2]. However, rodent control remains heavily reliant on rodenticide poisons - primarily anticoagulants [7] - which suffer significant drawbacks including severe welfare impacts on target species, bioaccumulation and exposure risks to non-target species, and high fixed operational costs [8,9]. These limitations underscore the urgent need for development of more precise and scalable tools for rodent control [10].

Genetic biocontrol of rodent populations offers a promising suite of species-specific suppression technologies with diverse mechanisms and control prospects. Although no genetic biocontrol tools for rodents are currently field-deployable, most development efforts to date have focused on self-sustaining population suppression systems which can, in principle, reduce rodent populations beginning from very small release sizes [11–13]. This feature is particularly important for rodents because individuals released as part of a biocontrol program may themselves contribute to predation. CRISPR-based homing population suppression gene drives have been explored in mice [14,15] and rats [16]. When targeting a gene required for female fertility or viability, these have potential to eliminate invasive rodent populations; however, to date their performance has been hindered by limited homing rates and frequent generation of drive-resistant alleles [14,15]. Modeling indicates that a modified design incorporating cleavage of an unlinked haplolethal gene could enable elimination even at currently-achievable homing rates [17]. Y-linked X chromosome shredders have also been explored in mice [18] due to their potential to bias inheritance while distorting the sex ratio towards males, eventually leading to population elimination. To date, significant male offspring-bias has not been achieved using this approach; however, modeling suggests the approach could be successful given improved efficiency parameters [19]. A conceptual variant in which a programmable endonuclease shreds the Y chromosome, converting males into fertile females, also shows promise based on preliminary molecular and modeling results [20]; but is yet to be developed. Finally, a gene drive has been modeled in mice that leverages the natural *t-*haplotype, which functions as a male meiotic drive system, and biases transmission of a linked CRISPR cassette targeting a female fertility gene [21]. In preliminary molecular work, this system has been engineered as a split-drive in mice, with encouraging estimates of biased inheritance and interruption of female fertility [21].

Each of the above-mentioned, self-sustaining tools is expected to increase in frequency following a small release, and to spread geographically. This is desirable when the goal is to eliminate invasive rodents on a wide scale with minimal releases and at low cost; but in the early phases of a biocontrol program, and depending on regulation and stakeholder engagement, it will likely be desirable to adopt a strategy that displays limited spread, both spatially and temporally, even if this requires continued releases during the control period. Strategies displaying this property are referred to as “self-limiting,” and have received relatively little attention for rodent control to date due to their large required release sizes. The most widely used self-limiting genetic biocontrol approach is the release of sterile males (SMR) [22–24]; but required release sizes are typically at least an order of magnitude larger than that of the target population [25]. This is appropriate for insect pest control, as it is invariably female insects that cause agricultural damage or transmit vector-borne diseases; but it renders the approach unsuitable for rodents, as both female and male rodents predate native species, with each release inflating the predator population. Several variations of SMR are available for insects, most notably fsRIDL (release of insects carrying a female-specific dominant lethal gene, in which the lethal gene persists in males and causes continued suppression in females for a few additional generations) [26]; however, release sizes remain high using this approach and may not be suitable for rodents.

A valuable addition to the rodent genetic biocontrol toolkit would be a self-limiting system capable of suppressing or eliminating populations with modest release sizes, but without the risk of substantial spread to neighboring areas. One such approach is the Y-linked genome editor (YLE), first proposed in a species-agnostic modeling study by Burt and Deredec in 2018 [27], and recently engineered in the malaria vector, *Anopheles gambiae* [28]. In this strategy, a nuclease cassette is inserted onto the Y chromosome and used to disrupt autosomal or X-linked genes essential for female viability or fertility [27]. Its strength lies in the fact that harmful effects only manifest in females, who do not have a Y chromosome, and hence the YLE is not inherently selected against. In the absence of a construct-based fitness cost, the YLE can therefore persist at its release frequency indefinitely, maintaining its suppressive effect, in contrast to other self-limiting strategies, such as SMR and fsRIDL, for which the population quickly recovers once releases are halted. That said; YLEs will be eliminated from a population in the presence of a construct-based fitness cost, at a rate proportional to the magnitude of the fitness cost, which could be engineered if desired [27]. Finally, because the YLE does not bias inheritance in its favor, it is not expected to increase in frequency following releases, and hence will not reach substantial frequencies in neighboring populations connected by migration. The path to engineering YLEs in rodents is promising, as integration and expression of Y-linked transgenes has already been achieved in mice [29], and several genes involved in female fertility and viability have already been identified [30,31].

Here, we use mathematical modeling to explore the potential and logistic requirements of YLEs as a self-limiting genetic biocontrol strategy for suppression of the invasive house mouse, *Mus musculus*. We use MouseGD, a stochastic modeling framework that incorporates inheritance patterns and rodent-specific life history [32], to simulate YLE release schemes on a small island setting as a case study. Small islands are often proposed as field sites for genetic biocontrol tools given their isolation and potential for proof-of-concept testing. We explore key product-related and operational parameters, such as the size and duration of releases, YLE-carrier fitness, and characteristics of the targeted female fertility/viability gene(s), to confirm that construct and release requirements are both favorable and attainable. We also benchmark YLEs against other self-limiting tools (SMR and fsRIDL), with emphasis on their release requirements, and self-sustaining tools (homing gene drives and X chromosome shredders), with emphasis on their current status and potential for confinement. Through this comparative, species-specific analysis, we confirm YLEs as the most efficient population suppression system for rodents that does not present a risk of invasive spread beyond its release site, highlighting the relevance of YLEs to the early phases of a rodent genetic biocontrol program, and possibly beyond.

## 2. Results

### Baseline dynamics of a Y-linked editor in an island mouse population

We evaluated the population-level impact of the YLE strategy on rodents using the MouseGD framework [32] parameterized for an isolated, randomly-mixing island setting with a carrying capacity, *K*, of 10,000 mice. A setting of this type could form the basis of an initial field release. The genetic mechanism modeled (**Fig 1a**) involves a YLE carried by males, targeting a critical, autosomal, female-specific gene during spermatogenesis. All male offspring of YLE-carrying males inherit the YLE and remain viable, while female offspring inherit a dominant, non-viable or sterile mutation, facilitating population suppression. Full details of the modeling framework and its parameterization are provided in the Methods section and **Tables A and B** in **S1 Text**, respectively.

**Fig 1.**
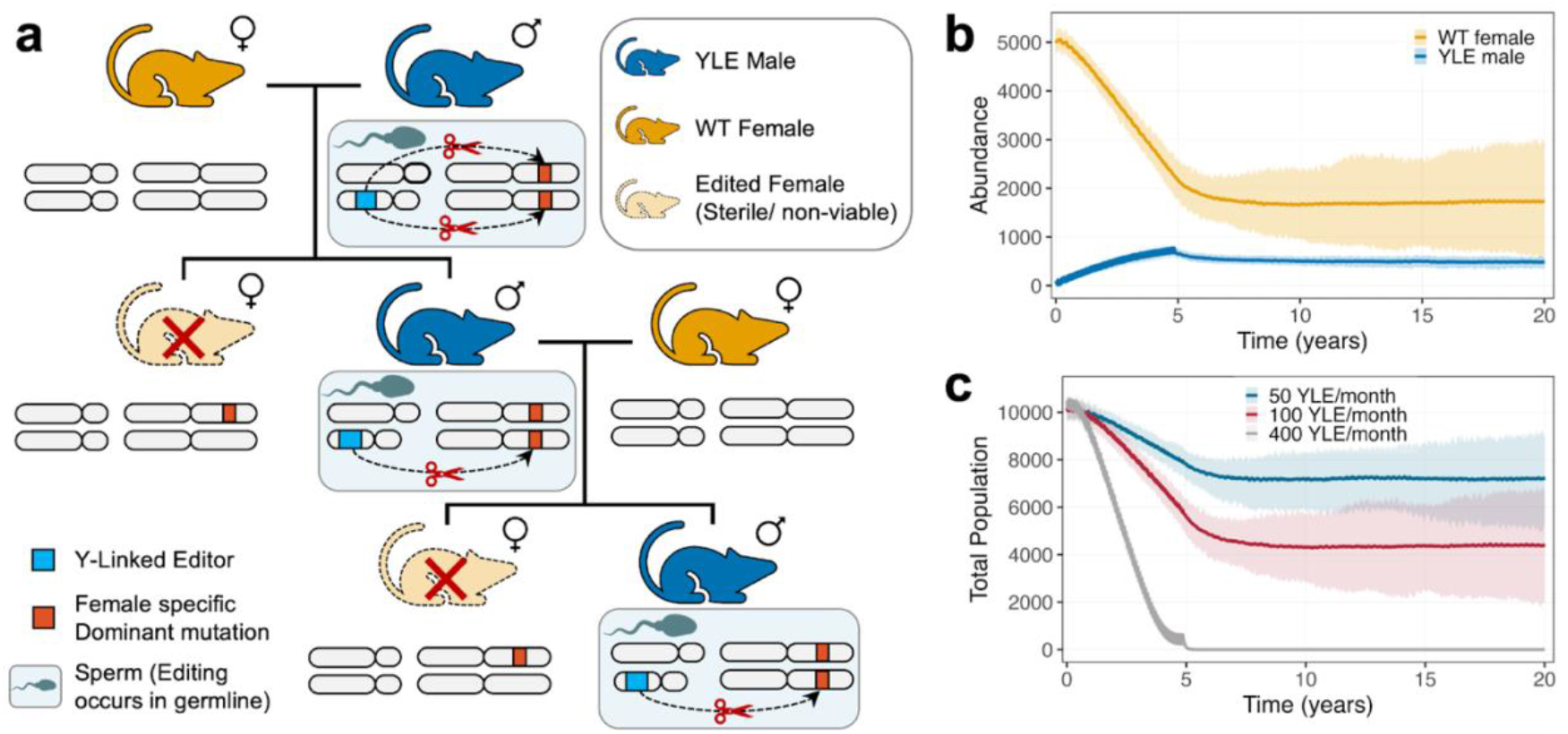
Y-linked editor (YLE) mechanism and population impact on house mice. **(a)** Schematic diagram of the inheritance pattern of a YLE (in blue) in mice. YLE-carrying males mate with wild-type (WT) females (in orange) producing offspring in which an autosomal gene required for female fertility or viability is edited (an X-linked gene may also be edited, but is not depicted). Female offspring that acquire the dominant mutation (depicted at the target site in red) are rendered either unviable or sterile (depicted as light orange with a dotted outline), while male YLE-carrying offspring survive and transmit the YLE to the next generation. **(b)** Population trajectories for stochastic simulations of a YLE release scenario in which 100 YLE males are released monthly for five years (equivalent to 2% of the initial adult male population released monthly). The YLE confers no fitness cost to male carriers and targets a gene required for female fertility. Total population counts are depicted in grey, and YLE male counts are depicted in blue. Solid lines depict mean population trajectories, and shaded ribbons represent 95% confidence intervals across 100 stochastic simulation replicates. **(c)** Population trajectories for release scenarios in which 50, 100 and 400 YLE-carrying males are released monthly for five years (equivalent to 1%, 2% and 8% of the initial adult male population released monthly). The full set of model parameters are provided in **Tables A and B** in **S1 Text**.

Baseline simulations of releases of YLE-carrying males reveal expected intervention outcomes and their dependence on release size (**Fig 1b-c**). Given a YLE that does not confer a male fitness cost, a moderate, five-year release scheme consisting of 100 released males per month (i.e., 2% of the initial male population per month) fails to achieve elimination; but establishes the YLE at a frequency of ~22% in the male population, and through mutation of a female sterility gene, maintains a continued suppression of the total mouse population of ~56% (**Fig 1b**). In contrast, a larger release scheme consisting of 400 released males per month (i.e., 8% of the initial male population per month) is successful at eliminating the island mouse population in 100% of stochastic simulation replicates, demonstrating that improved outcomes may be achieved at higher release sizes (**Fig 1c**). Of note, as explored in the next section, the level of suppression achieved is dependent on YLE-associated male fitness costs, and in the presence of such a fitness cost (whether manifest as a reduction in lifespan or mating competitiveness of YLE-carrying males), the YLE will eventually be eliminated from the population, which will recover to its equilibrium size.

### YLEs can eliminate an island mouse population within five years

To assess the viability of the YLE approach at eliminating an island mouse population, we systematically explored intervention outcomes while varying two model parameters that these outcomes are most sensitive to: i) the fitness cost of the YLE construct on male carriers (manifest as a reduction in male lifespan), and ii) the size of monthly releases. Outcomes considered are: i) the probability of elimination (PoE), i.e., the proportion of 100 stochastic simulations in which rodent elimination is realized, ii) the time to elimination (TTE), i.e., the median time by which elimination was achieved for scenarios in which the PoE is <0.5, and iii) the mean mouse population size measured over a 50-year period following the beginning of releases. In all simulations, we considered monthly releases conducted over a ten-year period, and a YLE targeting a gene required for female fertility, as we assessed this to be easier to engineer. Unless otherwise specified, default parameter values were used as specified in **Table B** in **S1 Text**. The sensitivity analysis that informed our choice of parameters to vary is provided in **Figs A and B** and **Table C** in **S1 Text**.

Results of these simulations, depicted in **Fig 2**, are encouraging. Eliminating the mouse population within five years is a feasible goal (dashed orange line in **Fig 2b**), requiring moderate release sizes (e.g., ≥7% of the initial male population per month) and moderate-to-small reductions in YLE-carrier fitness (e.g., male lifespan reductions ≤40%). Eliminating the mouse population within ten years can be achieved over a much wider range of parameter values (**Fig 2a**). A trade-off is evident between YLE-associated fitness costs and required release size. E.g., for a 20% lifespan reduction, monthly releases of 3% of the initial male population are sufficient to achieve a PoE ≥90% within ten years; while for an 80% lifespan reduction, monthly releases must be ≥5% of the initial male population to achieve the same result (**Fig 2a**). The same trade-off is seen for TTE, including for a target TTE within five years (**Fig 2b**). Where population elimination is not achieved, long-term population suppression is still observed, especially for YLE-associated lifespan reductions ≤30% (**Fig 2c**). These outcomes are sensitive to campaign design, with a YLE targeting a female-specific sterility gene offering slightly improved performance over one targeting a female-specific viability gene (**Fig C** in **S1 Text**), perhaps due to increased density-dependent deaths as sterile females remain in the population. Shortening the release period from ten to five years contracts the parameter space for success (**Fig D** in **S1 Text**).

**Fig 2.**
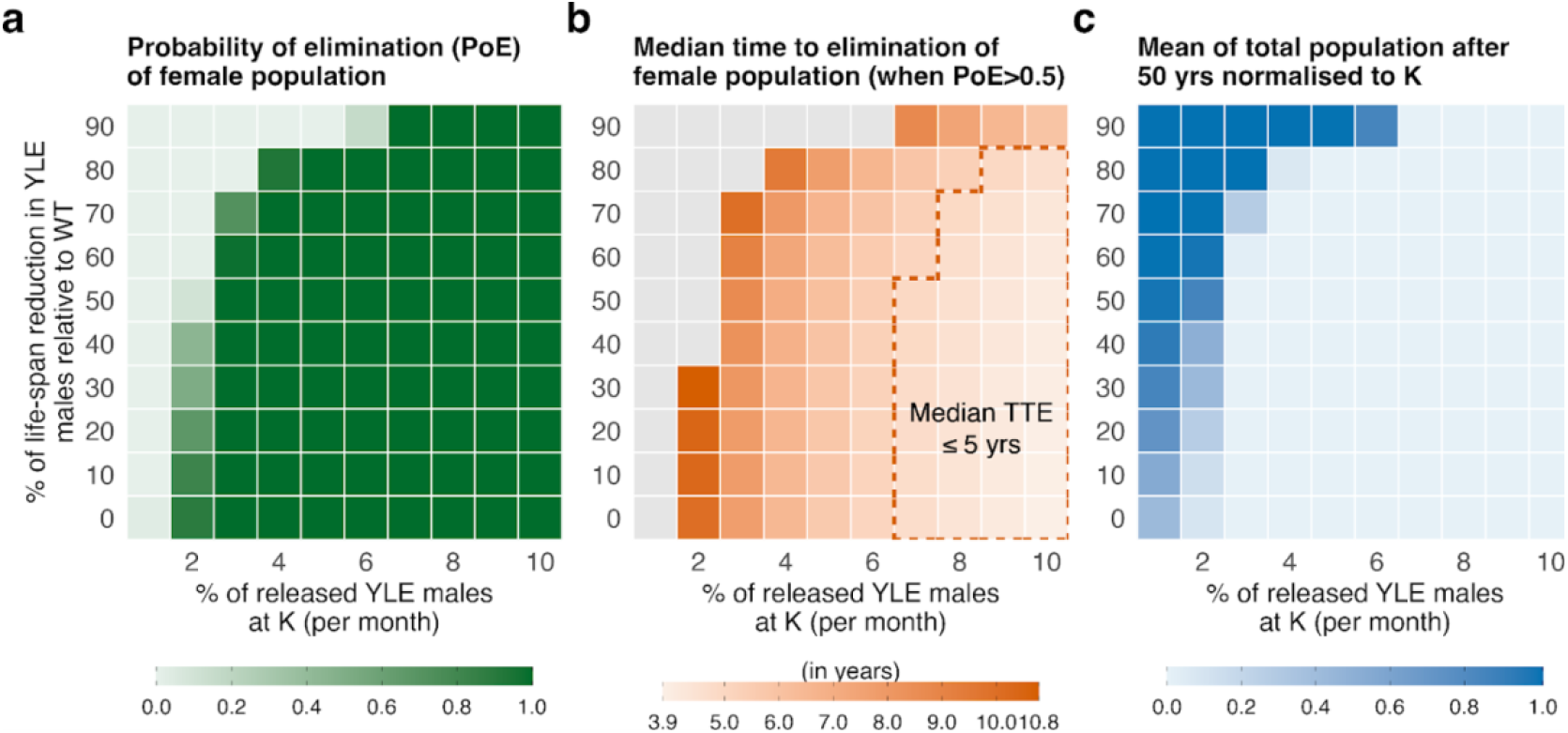
Simulated outcomes of Y-linked editor (YLE) release schemes as a function of the fitness cost to YLE-carriers and release size. Monthly YLE releases are simulated for a period of ten years in a randomly-mixing population of 10,000 adult mice at equilibrium. Three outcomes are tracked, each summarizing the results of 100 stochastic simulations: **(a)** the probability that the mouse population is eliminated, as measured by the fraction of simulations in which wild-type (WT) female mice are eliminated (darker green indicates a higher elimination probability); **(b)** the median time to elimination of WT females (in years) for cases where the elimination probability is ≥0.5 (darker orange indicates a higher time to elimination, grey represents an elimination probability <0.5, and the dashed orange perimeter marks the region of parameter space where the median time to elimination is ≤5 years); and **(c)** the mean total mouse population size for the 50-year period beginning with the first YLE release (here, the mean population is normalized with respect to the initial adult population size, with darker blue indicating larger population size). Outcomes are tracked for two parameters that model outcomes are most sensitive to: on the x-axis, the release size (measured as a percent of the initial adult male population size); and on the y-axis, YLE-carrier fitness (measured as a percent reduction in male lifespan). The full set of model parameters are provided in **Tables A and B** in **S1 Text**.

### Population elimination can be achieved with haplosufficient target genes for a more restrictive parameter space

The scenario of a haploinsufficient female-sterile/lethal target gene is ideal; however, identifying such a target gene and engineering gRNAs to disrupt it may prove difficult. We therefore evaluated the ability of a YLE targeting a haplosufficient or partially haplosufficient target gene to eliminate a small island mouse population (**Fig 3**). First, we considered a “worst case” scenario in which the YLE targets a fully haplosufficient recessive gene (i.e., only homozygous-edited females are non-viable) (**Fig 3a-c**). As anticipated, this strategy is significantly less efficient; but can still affect population suppression or elimination under permissive parameter values - required release sizes are roughly double those for a haploinsufficient target gene. E.g., for a 20% lifespan reduction, monthly releases ≥3% of the initial male population are required to achieve a PoE ≥90% within ten years for the haploinsufficient case, while monthly releases ≥6– 8% are required for the haplosufficient case. Population elimination within five years can also be achieved with a haplosufficient target gene; but monthly releases must be ≥18-20% of the initial male population and the lifespan reduction fitness cost must be ≤30% (**Fig 3b**) - a significantly more restrictive scenario.

**Fig 3.**
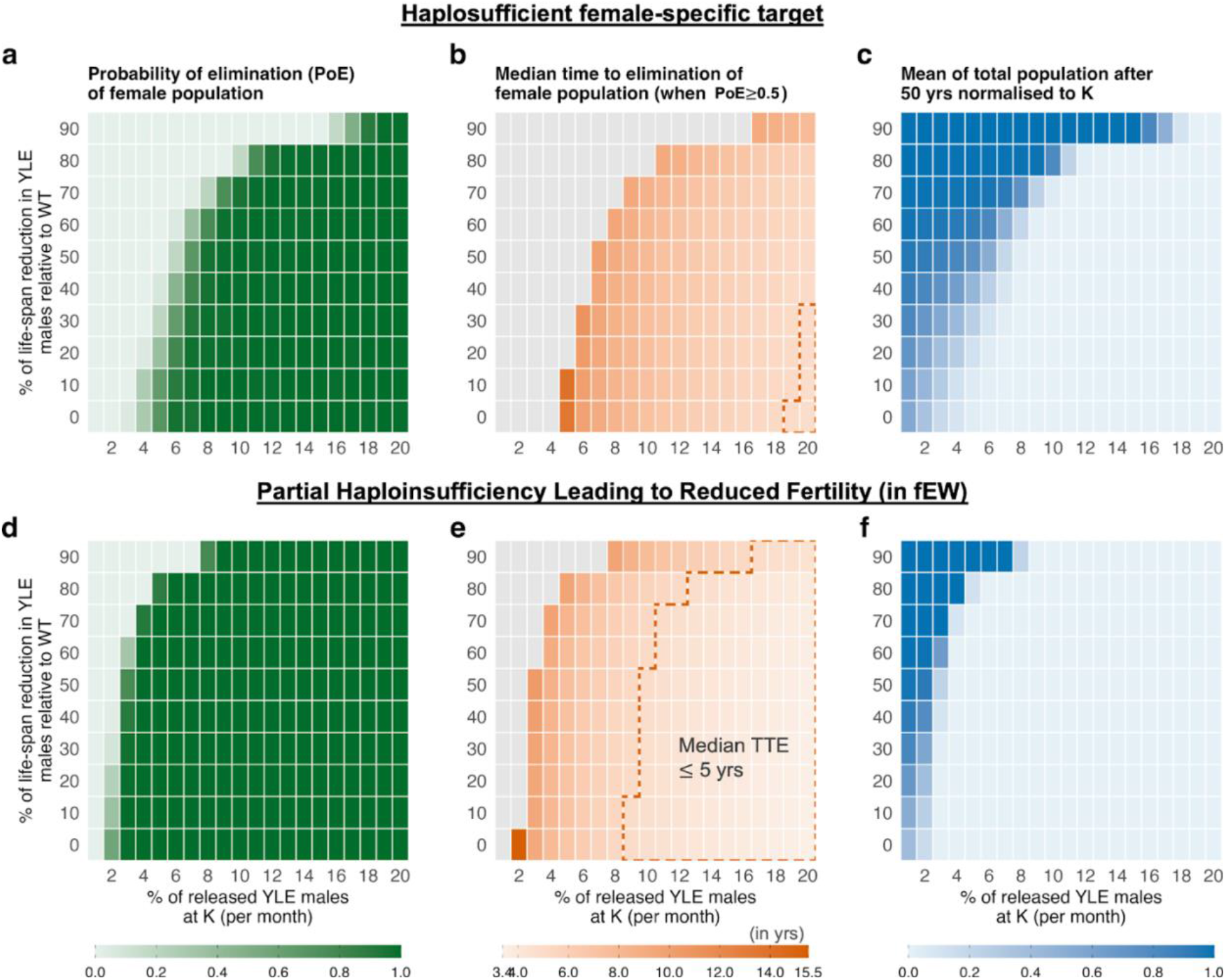
Simulated performance of Y-linked editors (YLEs) with haplosufficient and partial-haploinsufficient target genes as a function of fitness cost and release size. In panels **(a–c)**, the YLE targets a haplosufficient female-specific recessive fertility gene (i.e., females are only infertile when both gene copies are edited); and in panels **(d–f)**, the YLE targets a partial-haploinsufficient female-specific fertility gene (i.e., when one gene copy is edited, the heterozygous carriers have a 50% reduction in fertility, and when both gene copies are edited, they are infertile). Monthly YLE releases are simulated for a period of ten years in a randomly-mixing population of 10,000 adult mice at equilibrium. Three outcomes are tracked: **(a**,**d)** the probability that the mouse population is eliminated (darker green indicates a higher elimination probability); **(b**,**e)** the median time to elimination of wild-type females (in years, darker orange indicates a higher time to elimination, and the dashed orange perimeter marks the region of parameter space where the median time to elimination is ≤5 years); and **(c**,**f)** the mean total mouse population size for the 50-year period beginning with the first YLE release (here, the mean population is normalized with respect to the initial adult population size, with darker blue indicating larger population size). Outcomes are tracked for release size on the x-axis (measured as a percent of the initial adult male population size); and YLE-carrier fitness on the y-axis (measured as a percent reduction in male lifespan). The full set of model parameters are provided in **Tables A and B** in **S1 Text**.

Next, we investigated the case of a “partial-haploinsufficient” target gene, for which female fertility was reduced by 50% in heterozygotes, and 100% in homozygotes for the edited gene. YLE performance was significantly better than for the case of a haplosufficient target gene, in terms of both PoE and TTE (**Fig 3d-f**), and closely approached the performance of the ideal haploinsufficient benchmark. This is a useful realization for YLE development, as it suggests that partial-haploinsufficient target genes may be sufficient to achieve elimination of a mouse population within five years for attainable release sizes and YLE-carrier fitness costs. Thus, if such target genes are easier to identify, they are promising options for YLE-based rodent control.

### YLEs are more efficient than other self-limiting genetic biocontrol tools

Given that YLEs are classified as a self-limiting genetic biocontrol tool, we benchmarked their performance against two other such tools: i) SMR, the most commonly-used approach for insect pests; and ii) release of rodents carrying a female-specific dominant lethal gene (fsRRDL) - a rodent equivalent of insect-based method, fsRIDL. We focused our benchmarking against fsRRDL, as this is a more efficient approach than SMR - the lethal gene persists in males and causes continued suppression in females for a few additional generations before being eliminated due to its inherent fitness load in females. Simulations for a variety of monthly release sizes and male fsRRDL-carrier fitness costs reveal that this approach requires substantially higher releases to achieve the level of suppression or elimination achieved with releases of YLEs (**Fig 4a-c**). E.g., to guarantee elimination within ten years, fsRRDL requires monthly releases of ≥14% of the initial male population even in the absence of a male fsRRDL-carrier fitness cost. In comparison, a YLE with a haploinsufficient target gene can achieve similar outcomes with monthly releases of ≥3% of the initial male population (**Fig 2**), and a YLE with a haplosufficient target gene (the weakest YLE) can achieve this with monthly releases of ≥8% of the initial male population (**Fig 3a-c**).

**Fig 4.**
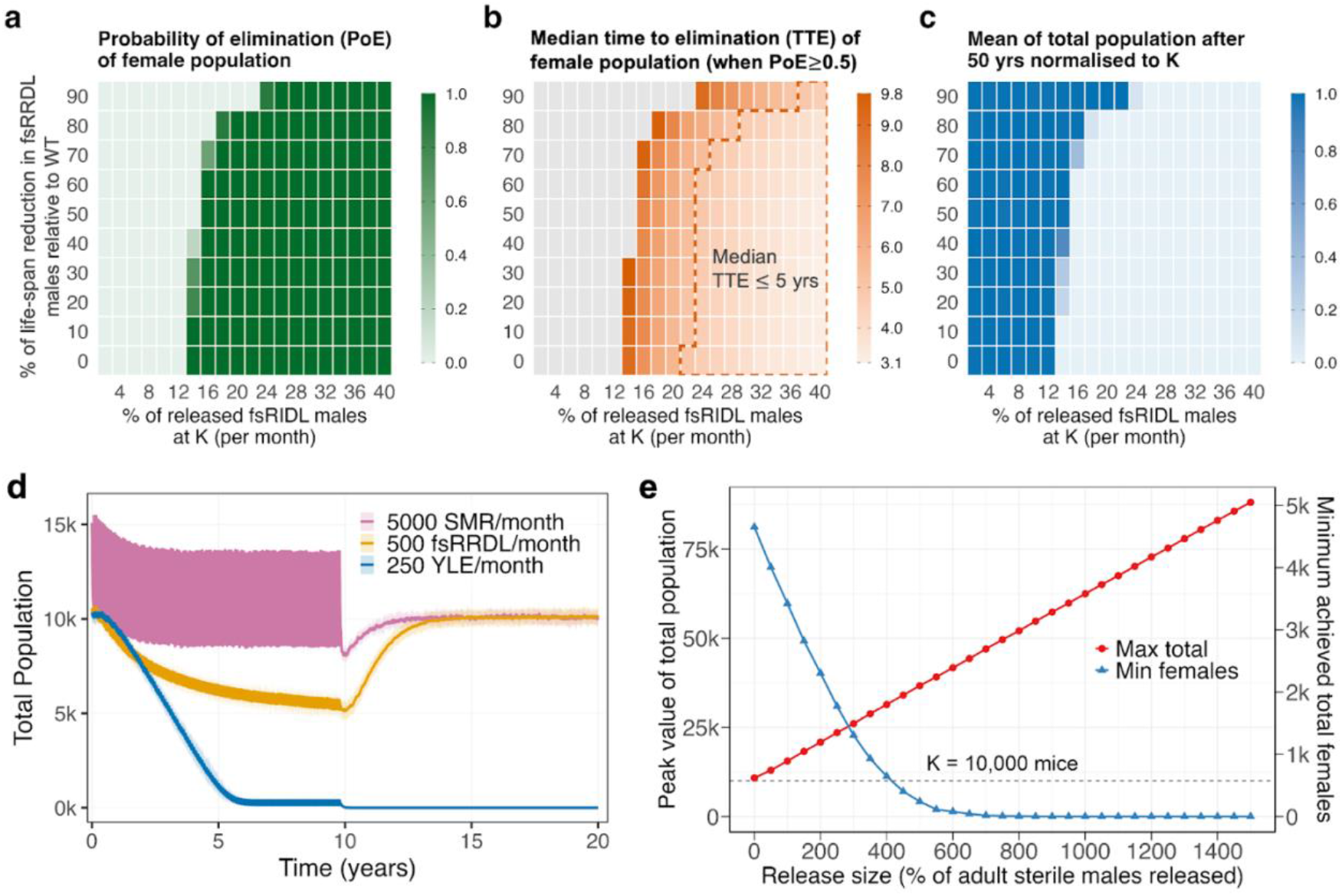
Comparative performance of Y-linked editors (YLEs), fsRRDL (release of rodents carrying a female-specific dominant lethal gene), and sterile male releases (SMR). **(a-c)** Monthly fsRRDL releases are simulated for a period of ten years in a randomly-mixing population of 10,000 adult mice at equilibrium. Three outcomes are tracked: **(a)** the probability that the mouse population is eliminated (darker green indicates a higher elimination probability); **(b)** the median time to elimination of wild-type females (in years, darker orange indicates a higher time to elimination, and the dashed orange perimeter marks the region of parameter space where the median time to elimination is ≤5 years); and **(c)** the mean total mouse population size for the 50-year period beginning with the first fsRRDL release (here, the mean population is normalized with respect to the initial adult population size, with darker blue indicating larger population size). Outcomes are tracked for release size on the x-axis (measured as a percent of the initial adult male population size); and fsRRDL-carrier fitness on the y-axis (measured as a percent reduction in male lifespan). **(d)** Total population trajectories for ten-year release campaigns of SMR (5,000 males released per month), fsRRDL (500 males released per month), and YLE (haploinsufficient target gene, 250 males released per month). Mean population trajectories are represented by solid lines, and 95% confidence intervals are represented by a shaded light color. **(e)** Mean of peak total population (red) and minimum WT female population (blue) achieved during a ten-year SMR campaign as a function of monthly release size. The dashed dark grey line represents the equilibrium population size in the absence of interventions.

The efficiency gap between YLEs and other self-limiting genetic biocontrol tools (SMR and fsRRDL) is illustrated in direct trajectory comparisons in **Fig 4d**. Interventions using SMR must truly be inundative, highlighting their inapplicability for rodent suppression. Even at a 1:1 release ratio, which would approximately double the male rodent predator population temporarily, the intervention fails to achieve more than a 30% transient reduction in the female rodent predator population. The logistical barrier to utilizing this approach is highlighted in **Fig 4e**, which displays the peak total rodent population for a variety of release sizes. This reaches 16,000 for the SMR 1:1 release scenario (i.e., 160% of the initial rodent population size). To eliminate the rodent population within ten years using this approach, our model predicts that SMR releases would need to exceed 800% of the initial male rodent population size, causing the total mouse population to transiently swell to ~50,000 (i.e., five times its original size), and leading to significant increases in predation of native species during the control period.

The fsRRDL strategy is more efficient than SMR; but suffers the same issue that releases must be substantially larger than for YLEs. As depicted in the trajectory comparisons in **Fig 4d**, for a monthly release size of 500 fsRRDL males (i.e., 10% of the initial male rodent population per month), the intervention achieves a ~58% reduction in the female rodent predator population at the end of the ten-year release period, and the total rodent population reaches a maximum of ~11,000 (i.e., a 10% increase) (**Fig E** in **S1 Text**). Monthly fsRRDL male releases of ≥14% of the initial male population are predicted to lead to elimination in the absence of fsRRDL-carrier fitness costs (**Fig 4a**), causing the rodent predator population size to again transiently increase (**Fig E** in **S1 Text**). In contrast, YLEs are simulated to achieve population elimination for much smaller release sizes, which also means that any transient swell in rodent population size is less pronounced. Trajectory comparisons in **Fig 4d** depict the case of a YLE targeting a haploinsufficient female fertility gene. In the absence of a YLE-carrier fitness cost, this eliminates the rodent population within five years for a monthly release size of 250 males (i.e., 5% of the initial male population). There is a 3% rise in the total rodent predator population size; but this is more than compensated for by the substantial suppression during the control period (**Figs F and G** in **S1 Text**).

### YLEs can achieve similar elimination outcomes with better confinement than homing gene drives and X-shredders

In contrast to self-limiting strategies, which require continued releases, self-sustaining genetic biocontrol tools (i.e., gene drives) are designed to actively propagate through a population following a relatively small release. We modeled two prominent self–sustaining strategies proposed for implementation in rodent systems: i) a suppression homing gene drive (HGD), which uses a nuclease to convert WT alleles to drive alleles in heterozygotes, interrupting a critical female fertility gene [14–16]; and ii) a Y-linked X-shredder, which destroys the X chromosome during male gametogenesis to produce a heavily male-biased population, eventually leading to a population crash [18,20]. For both systems, adequate parameters (e.g., rates of homing or sex ratio distortion) are yet to be achieved. Therefore, we simulated two scenarios: i) an ideal scenario, in which homing or sex ratio distortion is perfect and there is no resistant allele formation; and ii) a non-ideal scenario, in which parameter values resemble those already achieved in the lab (**Tables D and E** in **S1 Text**). In agreement with prior modeling [17,19], simulation output in **Fig 5a** conveys that, while current implementations of these tools are expected to lead to transient suppression before resistant alleles appear and are selected for, inducing a population rebound, ideal versions of these systems are exceptionally powerful, eliminating rodent populations within ~3 years beginning from a single release of 250 transgenic males (equivalent to just one monthly release for the YLE strategy). In contrast, the YLE strategy reliably achieves elimination, albeit for a modest increase in duration of releases and elimination timeline (pictured in **Fig 5a** for monthly releases of 250 YLE-carrier males, achieving elimination within ~6 years).

**Fig 5.**
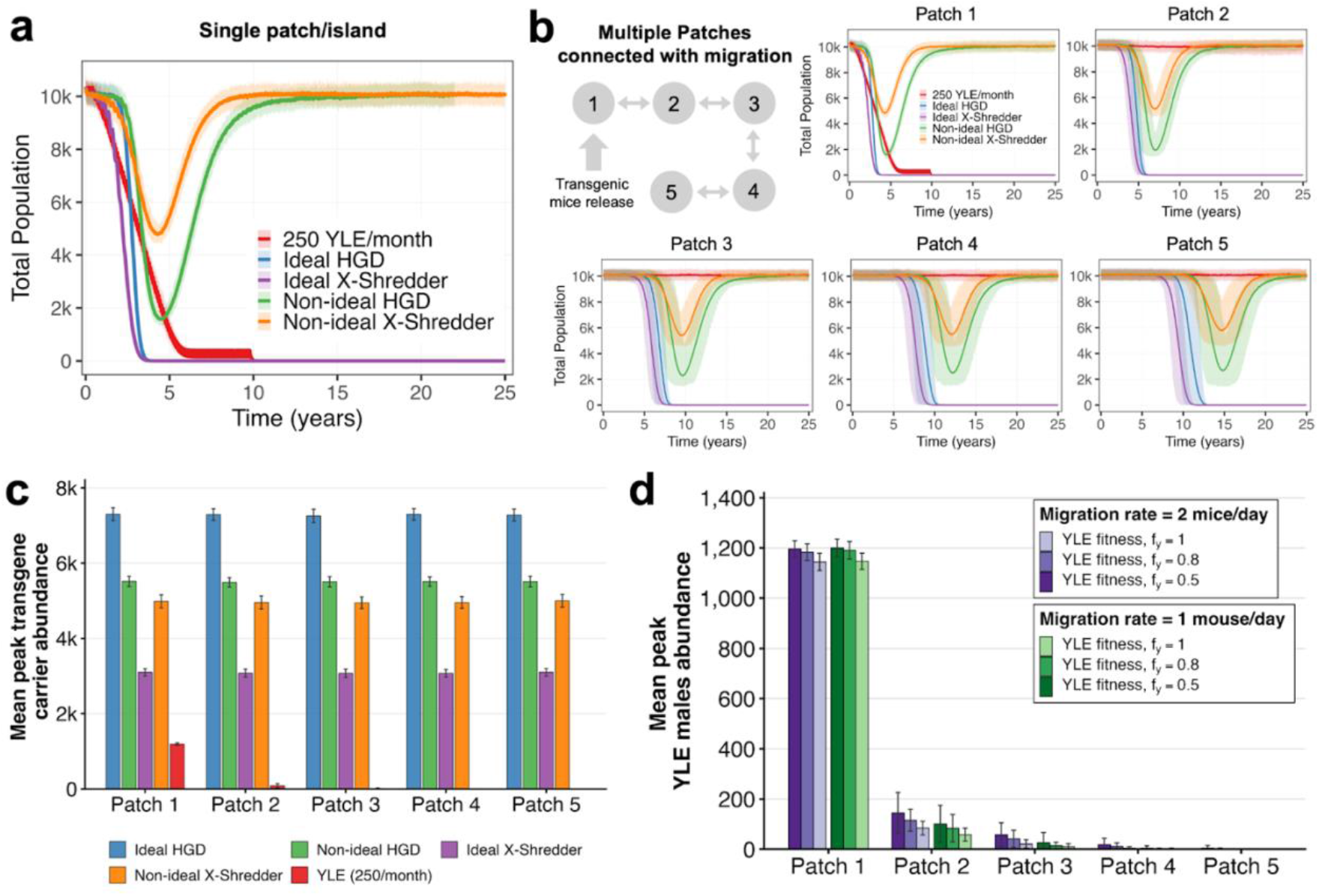
Comparative performance and confinability of self-sustaining gene drives and a Y-lined editor (YLE) strategy. The homing gene drive (HGD) and Y-linked X-shredder strategies involve a single release of 250 adult males, while the YLE strategy involves monthly releases of 250 males for ten years. Ideal drive scenarios assume perfect homing, sex ratio distortion, and no resistance formation. Non-ideal drive scenarios incorporate observed constraints such as high rates of resistant allele formation. Parameters for all systems (e.g., cutting rates, repair outcomes, and resistance formation) for both ideal and non-ideal scenarios are detailed in **Tables B, D and E** in **S1 Text. (a)** Population dynamics for five modeled systems (ideal and non-ideal HGDs and X-shredders, and YLEs) in a single isolated patch. Plots show the mean trajectory of total population size (solid lines) and the 95% confidence interval (lightly shaded region) across 100 stochastic replicates. **(b)** Spatiotemporal dynamics in a “stepping stone” metapopulation model. Patch 1 represents the target population receiving transgenic releases, patches 2-5 represent serially connected non-target populations (migration rate of one mouse per day). **(c)** Mean peak abundance of transgene-carriers (individuals carrying Cas9 or homing alleles) across the target and non-target patches and 25-year simulation timeline. **(d)** Mean peak abundance of YLE-carrier males across target and non-target patches under varying YLE-carrier fitness costs (lifespan reductions of 0, 20 and 50%), and migration rates (1-2 mice per day).

To compare the potential for spatial confinement of each of these tools, we extended our analysis to a five-patch metapopulation, where patch 1 represents the target island, and patches 2–5 represent connected non-target populations arranged as a series of linear “stepping stones” (**Fig 5b**). For a migration rate of one mouse per day, the self-sustaining strategies (HGDs and X-shredders) display invasive behavior, spreading across all five populations, even for non-ideal gene drive parameters. In contrast, the YLE strategy achieves elimination of the target patch, with minimal spread to non-target populations. We further quantified confinement by analyzing the mean peak abundance of transgene-carriers across the metapopulation for each strategy (**Fig 5c**). Importantly, the peak abundance of gene drive-carriers is high (~3,000-7,000) across the metapopulation, while the peak abundance of YLE-carriers rapidly decays with distance, nearing zero in patches 2–5. Lastly, we assessed the sensitivity of results on the confinability of YLE-carriers to variations in fitness costs (i.e., reductions in YLE-carrier lifespan) and migration rates (**Fig 5d**). As expected, increasing the migration rate to two mice per day led to a modest increase in the peak population of YLE-carriers in non-target patches. Consistent with a prior YLE modeling study [27], we found that fitness costs on YLE-carriers yield lower peak abundances in non-target patches, in addition to decreasing the duration of YLE persistence in these patches, hence enhancing their confinement potential.

## 3. Discussion

Our modeling analysis demonstrates that YLEs are a promising self-limiting genetic biocontrol strategy for suppressing or eliminating invasive rodent populations on islands. Using the MouseGD framework [32], we showed that YLEs - by selectively inducing sterility or unviability in female offspring while avoiding direct fitness costs to male carriers - can achieve substantial and sustained reductions in rodent population size. Under a broad parameter space, including modest monthly releases and achievable construct-associated fitness costs, YLEs were simulated to eliminate an island mouse population with an equilibrium population size of 10,000 within a five-year period. Rodent elimination was simulated for both haploinsufficient and haplosufficient target genes, with both being viable strategies; but larger release sizes being required for the (weaker) haplosufficient case. Together with the relative confinability of YLEs to their release site, our results highlight the suitability of YLEs for the early phases of a rodent elimination program, and possibly beyond.

A key finding of this study is that required YLE release sizes to achieve elimination are well within the means of modern rodent production facilities, and given their modest size, are not expected to significantly increase the rodent predator population during the initial weeks of an intervention. For the small island case study with an equilibrium population size of 10,000, and considering a 20% lifespan reduction for YLE-carriers, a monthly release size of ≥7% of the initial male population is capable of eliminating the rodent population within five years, while a monthly release size of ≥3% of the initial male population is expected to achieve elimination within ten years. These scenarios equate to releases of 150-350 YLE-carrying males per month. Applying the statistical framework of Milchevskaya *et al*. [33] for rodent group size planning, this equates to 58 breeding pairs to ensure the availability of at least 150 male pups per month with 90% confidence, and 130 breeding pairs to produce 350 male pups per month. This scale of production is well within existing capabilities - one of the first mouse production facilities in Germany in 1929 was capable of producing over 4,000 mice per month [34], and in the US, laboratories currently produce ~9 million rodents per month [35]. Indeed, it may be possible to scale-up YLE-carrier mouse production severalfold to achieve rodent control on a much larger scale. Encouragingly, our models predict that, for releases of this size, the total rodent predator population is only expected to increase by ~3% for a brief period after releases begin and before suppression takes hold.

Compared to other self-limiting genetic biocontrol tools, our simulations convey a distinct efficiency advantage of YLEs over alternatives - SMR and fsRRDL. SMR requires inundative releases, often exceeding the wild male population size by an order of magnitude, to achieve elimination. This is impractical and counterproductive for rodents as each released male also acts as a predator. Female-specific dominant lethal systems such as fsRRDL are significantly more efficient than SMR; but still require release sizes 2-3 times larger than those required for YLEs. Furthermore, SMR and fsRRDL produce transient suppression that rebounds quickly after releases cease, whereas YLEs maintain a persistent suppressive effect because the Y-linked construct persists, indefinitely in the absence of a YLE-carrier fitness cost, and for an extended period in the presence of a modest fitness cost. This underscores an important advantage of YLEs over other self-limiting tools - suppression achieved using YLEs is cumulative, always building upon that achieved through prior releases.

Benchmarking YLEs against self-sustaining tools (i.e., gene drives) clarifies their niche within the rodent genetic biocontrol landscape. Ideal HGDs and X-shredders are extremely powerful, capable of eliminating populations from very small releases; but this comes at the cost of potentially uncontrollable spread to non-target populations. This may raise regulatory and stakeholder concerns, particularly in the early phases of a rodent control program. As our simulations demonstrate, even non-ideal gene drives have the potential to spread invasively. Furthermore, current molecular assemblies of gene drive systems in rodents remain limited by low homing rates, inadequate sex ratio distortion, and frequent resistant allele generation [14,15,18]. Engineering efforts seek to resolve these issues, for instance by improving temporal regulation of Cas9 during meiosis to increase conversion efficiency [14,36], and models of alternative drive designs suggest hopeful paths to rodent elimination given current parameter values [17]; however, issues regarding potentially uncontrollable spread to non-target populations remain. In contrast, YLEs do not induce an inheritance bias, and hence migration from the release site will not lead to invasive spread. Low levels of spillover to neighboring populations are expected; but these quickly decay with distance from the release site. If desired, confinement of YLEs can be enhanced by engineering a fitness cost on male YLE-carriers [27].

Our modeling necessarily relies on simplifying assumptions that introduce limitations. Most notably, we have neglected polyandry, which is prevalent in natural mouse populations - by some measures, nearly 50% of litters may be sired by multiple males [37]. Combined with sperm competition, this may reduce the reproductive success of modified males [37], potentially decreasing the impact of a YLE and requiring larger release sizes to achieve similar outcomes. We have assumed YLE-associated fitness costs to reduce male lifespan, rather than reducing male mating competitiveness. Sensitivity analyses could determine the implications of this assumption, although males will obtain fewer mates in both cases. We have assumed random mixing, which may be suitable for the small island setting considered here; but will break down for larger islands and landmasses, where spatial structure and barriers to dispersal shape mating networks. Regarding the inheritance component of the model, we have neglected resistant alleles in the female essential gene. Such alleles, which may be formed via NHEJ, could prevent cleavage of this gene, and if they maintain its function, would be selected for over cleavage-susceptible alleles in the presence of a YLE. While subsequent modeling efforts should address this concern, engineering solutions are available, such as selecting highly-constrained target sites, multiplexing of gRNAs, or targeting multiple female essential genes to further mitigate risk. While our modeling analysis has focused on autosomal target genes, targeting X-linked female-essential genes is also possible. We justify this choice on the basis that Burt & Deredec [27] found YLEs perform similarly whether the target gene is on an autosome or X chromosome, and autosomal targets may be easier to identify and validate. Finally, we have not considered seasonal effects on rodent populations in this analysis. This is within the scope of our modeling framework [38,39], and could potentially increase the chances of elimination during seasonal lows in natural population size.

From a practical engineering perspective, constructing a YLE in mice or rats appears within reach of current genome engineering capabilities. This would involve the targeted insertion of a nuclease cassette into a transcriptionally-active region of the Y chromosome, coupled with germline-specific expression to ensure cleavage of a female-essential gene (either haploinsufficient or haplosufficient) during spermatogenesis, or in the zygote immediately after fertilization via paternal deposition. In mice, the feasibility of sex chromosome genome engineering has been demonstrated by Douglas *et al*. [29], who expressed the Cas9 nuclease from sex chromosome-linked loci to generate single-sex litters and sex-specific phenotypes, underscoring the practicality of Y-linked nuclease deployment in mammalian systems [29]. To build upon this foundational work, a YLE could be engineered to target an autosomal or X-linked female-essential gene, where loss of function in daughters leads to sterility or unviability, but heterozygous sons remain unaffected. Potentially suitable target genes include *H1foo* (an oocyte-specific histone variant required for early embryogenesis) [40], *Gdf9* and *Bmp15* (oocyte-secreted growth factors essential for folliculogenesis) [41], *Zp1/Zp2/Zp3* (zona pellucida components required for fertilization) [42], *Mater/Nlrp5* (maternal-effect genes controlling embryonic cleavage) [43], and many other potential targets [31].

Efficient genome editing would rely on generating high rates of biallelic cleavage in edited sperm, which is achievable because nuclease protein and gRNAs can cleave during spermatogenesis prior to fertilization. Because the Y chromosome is inherited strictly paternally, males carrying the YLE experience no direct fitness cost from disruption of female-essential genes, enabling the construct to maintain its frequency in the population unless carrying an associated male fitness burden [27]. Collectively, the availability of a characterized Y-linked Cas9 that supports expression during spermatogenesis [29], and numerous validated female-essential target genes, makes construction of a YLE in mice feasible using current genome engineering technologies.

Together, these findings position YLEs as a compelling addition to the rodent genetic biocontrol toolkit. Their combination of efficiency, confinement, modest release requirements, and technical feasibility makes them well-suited for early-phase field deployment on small islands - settings where risk, reversibility, and regulatory oversight are paramount. While reinvasion risk remains a challenge common to all suppression tools, necessitating long-term biosecurity and monitoring, YLEs could play a transformative role in phased elimination strategies, serving either as a standalone solution, or as a complement to other genetic or conventional control methods. As molecular improvements enable precise Y chromosome engineering and address fitness costs and resistant allele formation, YLEs may ultimately emerge as a scalable, ecologically-responsible, and publicly-acceptable approach to invasive rodent control and native species preservation.

## 4. Methods

Simulations for this study were carried out using the MouseGD framework [32], a rodent-adapted version of the mosquito-centric MGDrivE framework [44]. Like MGDrivE, MouseGD includes modules for inheritance, life history and landscape. The inheritance module describes the distribution of offspring genotypes given maternal and paternal genotypes, and is shared across the two frameworks. Inheritance patterns for SIT/SMR, fsRIDL/fsRDDL, Y-linked X chromosome shredders, and population suppression homing gene drives are included in MGDrivE [44]. The inheritance pattern of YLEs was incorporated into the framework as part of this study, and is detailed below. The life history module describes the development of mice through four life stages - gestating, nursing, adolescence and adult - and is elaborated upon below. We assumed a single, randomly-mixing population in this study, due to the small size of the modeled island population, and so did not incorporate spatial structure in the landscape module.

### Mouse life history and demography

In this study, we modeled the house mouse, *Mus musculus*. Life history for this species is divided into four stages: gestating (19 days), nursing (23 days), adolescence (37 days), and adult (690 days). Individuals move through these stages with daily transition probabilities based on the average duration of each stage (**Fig 6a**). Daily mortality is applied at the adult stage, with a density-independent rate, *μ*_*Ad*_, and a density-dependent rate that is a function of the environmental carrying capacity for adults, *K*, and a shape parameter, *Θ*. Reproduction follows mouse-specific biology whereby each adult female has a daily probability of conception, and the number of newly pregnant females follows a binomial distribution such that only a subset of the total female population is pregnant at a given time. The litter size of each pregnancy is drawn from a Poisson distribution with mean of *Q*. Life history parameters, descriptions and values are provided in **Table A** in **S1 Text** and described illustratively in **Fig 6a**.

**Fig 6.**
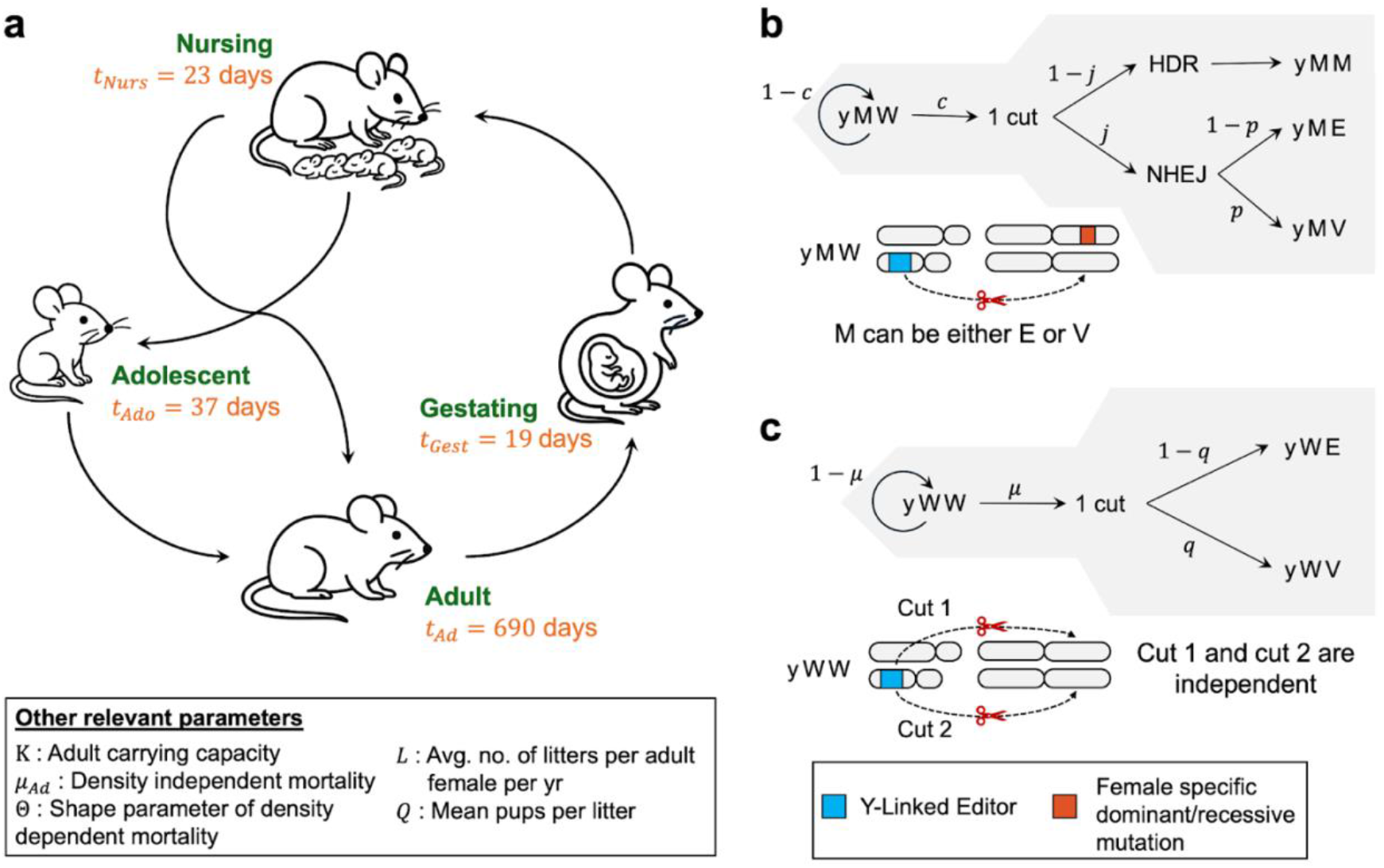
Mouse life history and Y-linked editor (YLE) inheritance dynamics used in MouseGD. **(a)** Schematic of the female life history, depicting transitions between nursing, adolescent, gestating and adult life stages, with associated mean stage durations and other demographic parameters listed in the legend. **(b)** Conversion dynamics in heterozygous males (yMW) carrying the YLE (y), one wild-type target allele (W), and one edited allele (M) at an autosomal locus. Of note, M represents either a dominant lethal mutation, E, or a recessively lethal variant, V. The target allele, W, is cleaved with probability, *c*. Following a cut, homology-directed repair (HDR) occurs with probability, (1−*j*), and non-homologous end-joining (NHEJ) occurs with probability, *j*. Conditional on NHEJ, the outcome is an edited allele, E, with probability, (1−*p*), or a resistant allele, V, with probability, *p*. **(c)** Conversion dynamics in homozygous wild-type males at the autosomal locus (yWW), where the YLE cuts each wild-type target independently. A cut occurs with probability, *μ*, at each target site, and repair yields an edited, E, or variant, V, allele with probabilities, (1−*q*), and *q*, respectively.

### Y-linked editor inheritance dynamics

The YLE was implemented in MouseGD [32] by specifying its genetic inheritance through an inheritance cube, a structure within the original version of MGDrivE [44] that defines the probabilities of offspring genotypes for all possible matings together with genotype-specific fitness parameters. Male genotypes are described by the type of Y chromosome they carry, either wild-type (Y) or edited/carrying the YLE construct (y), along with two autosomal alleles at the target site (W for wild-type, E for the desired mutation/edit, and V for a variant mutation). Female genotypes carry two X chromosomes and the same autosomal locus. Collectively, this leads to a total of 18 possible genotypes (six female and 12 male).

An important feature of the YLE is that a male with an edited Y chromosome (y) can modify wild-type autosomal alleles during gamete formation. In the YLE inheritance model, autosomal alleles are denoted as W (wild-type), E (desired mutant allele), and V (variant resistant mutant allele), while sex chromosomes are represented as X for females, Y for wild-type males, and y for males carrying the YLE construct. Editing occurs only in males carrying the YLE, with effects restricted to autosomal W alleles. In yWW homozygotes, each W allele is cleaved with probability, *µ*, and repaired to either E with probability, (1−*q*), or V with probability, *q*; otherwise, the allele remains unmodified (**Fig 6c**). In heterozygotes (yWE or yWV), the wild-type allele is cleaved with probability, *c*. Cleaved alleles are repaired by homology-directed repair (HDR) with probability, (1−*j*), converting the W allele to match the homologous allele (E in yWE, or V in yWV), or by NHEJ with probability, *j*, producing E with probability, (1−*p*), and V with probability, *p* (**Fig 6b**). If cleavage does not occur, with probability, (1−*c*), the wild-type allele is inherited. Males lacking the YLE and all females inherit alleles according to Mendelian segregation without modification. Because the editor is Y-linked, no cleavage occurs in females.

To explore the sensitivity of the YLE strategy to the underlying biology of the target locus, the YLE inheritance cube incorporates genotype-specific fitness parameters for three distinct genetic architectures: haploinsufficiency, haplosufficiency, and partial haploinsufficiency. We primarily investigated these architectures in the context of female fertility; however, to explore robustness, we also modeled effects on female viability (results detailed in **S1 Text**). For a haploinsufficient target, the edited allele (E) was modeled as dominant; consequently, any female carrying an edited allele (fEW, fEE, fEV) was assigned zero fitness (fully sterile or non-viable). In this scenario, the non-functional resistance allele (V) acts as recessive: homozygotes (fVV) have zero fitness, while heterozygotes retaining one wild-type copy (fVW) maintain full fitness. In contrast, a haplosufficient target was modeled as fully recessive, where a single WT allele preserves function. Here, all heterozygotes (fEW, fVW) retain full fitness, and only females lacking a functional WT copy (fEE, fEV, fVV) are rendered sterile (or non-viable). Finally, to represent partial haploinsufficiency, we modeled a scenario of incomplete dominance where heterozygous females (fEW) experience a 50% reduction in fitness, while females lacking a functional WT copy (fEE, fEV, fVV) have zero fitness. In all scenarios, the release males were specified as yEE, representing individuals carrying the Y-linked editor and homozygous for the edited allele at the autosomal target locus.

### Simulation scenarios and release designs

All simulations were run using daily time steps for up to 50 years from the time of the first release. Populations were initialized at carrying capacity and allowed to run for eight years before the release started, as a burn-in period to let the life stages reach a stable distribution. After this, gene drive introduction was modeled by releasing YLE males, typically for ten years unless otherwise mentioned, and in specified release proportions of the equilibrium male adult population. Simulations were carried out in R using MouseGD [32].

### Sensitivity analysis

We used a global sensitivity analysis to quantify how model parameters related to YLE inheritance and release scheme influence suppression and elimination outcomes. This analysis employed Latin hypercube sampling and boosted regression trees, closely following the approach of [21]. First, we generated 10,000 unique parameter combinations over the ranges listed in **Table B** in **S1 Text** using Latin hypercube sampling. To maximize coverage while minimizing compute, we ran one stochastic simulation per combination. For each simulation, we recorded: i) whether elimination occurred, and ii) the time to elimination (TTE), as measured from the start of releases. Elimination was operationally defined as the total number of females falling below one at or before the simulation horizon. Runs that never crossed this threshold were labeled “unsuccessful” (even if the population stabilized at a lower equilibrium).

Before modeling, we optionally restricted analyses to prespecified subranges (**Table B** in **S1 Text**) and removed predictors with zero variance within the filtered data. All predictors were treated as numeric. We then fitted two boosted regression trees using dismo::gbm.step (which wraps gbm): i) the probability of elimination, based on a binomial (Bernoulli) family with response (y{0,1}) (unsuccessful/successful), and ii) the TTE, based on a Poisson family with the response as integer days to elimination (only successful runs contributed to this model, as per the study design). Both models used the same learning settings: tree complexity = 3, learning rate = 0.01, bag fraction = 0.75, and fivefold cross-validation. To summarize parameter importance, we extracted the relative influence (percentage contribution) from the cross-validated optimal number of trees for each boosted regression tree.

## Supporting information

**S1 Text**. Supplemental analyses, figures and tables.

## Data availability statement

All code and source files used to produce the figures and analysis in this manuscript are available at https://github.com/prateek1verma/YLE-Rodent-Control/. The code is available under the GPL3 License and is free for other groups to modify and extend as needed.

