## Supplementary material for "Y-linked editors for invasive rodent control": S1 Text

### S1 Text: Supplemental analyses, figures and tables

### Supplementary figures

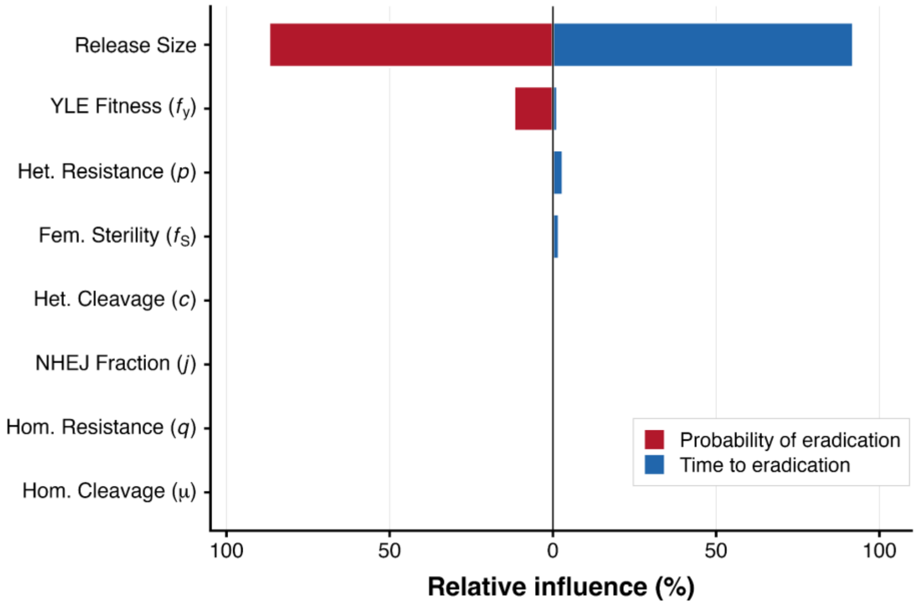


**Fig A. Relative influence of input parameters on eradication outcomes based on boosted regression tree (BRT) analysis.** The influence of seven model parameters on two distinct outcomes was evaluated: the likelihood of achieving successful eradication (probability of eradication) and, among successful simulations, the time required to reach eradication (time to eradication). Bar values represent the relative contribution (percentage) of each parameter to the predictive performance of the respective BRT models. The influence scores sum to 100% for each outcome separately. Parameters with larger values have a greater impact on the model's predictions for that specific outcome.

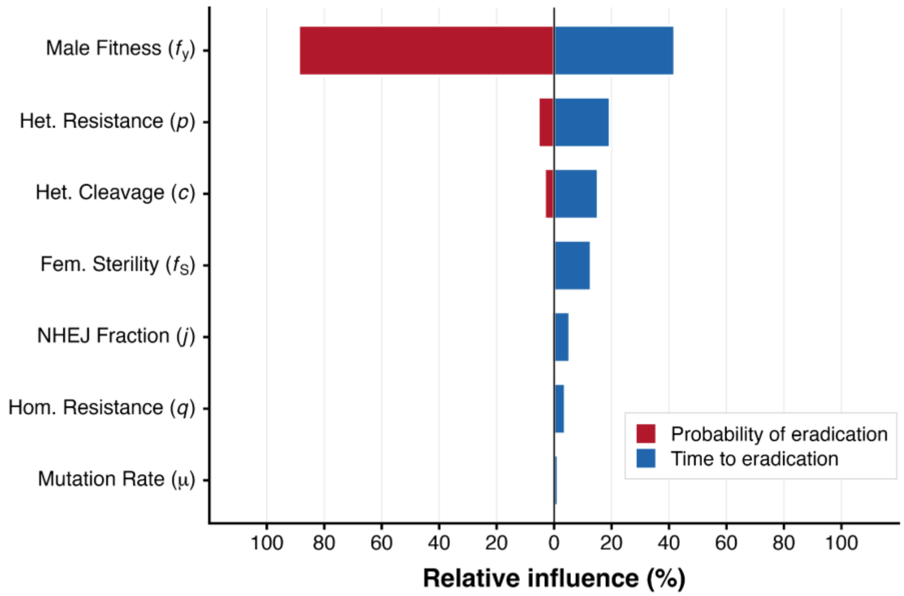


**Fig B. Relative influence of input parameters with fixed release size.** BRT analysis of parameter contributions to eradication probability and time to eradication. Here, the relative release size was held constant at 3% of adult male carrying capacity. All other parameter ranges and modeling definitions are identical to **Fig A**.

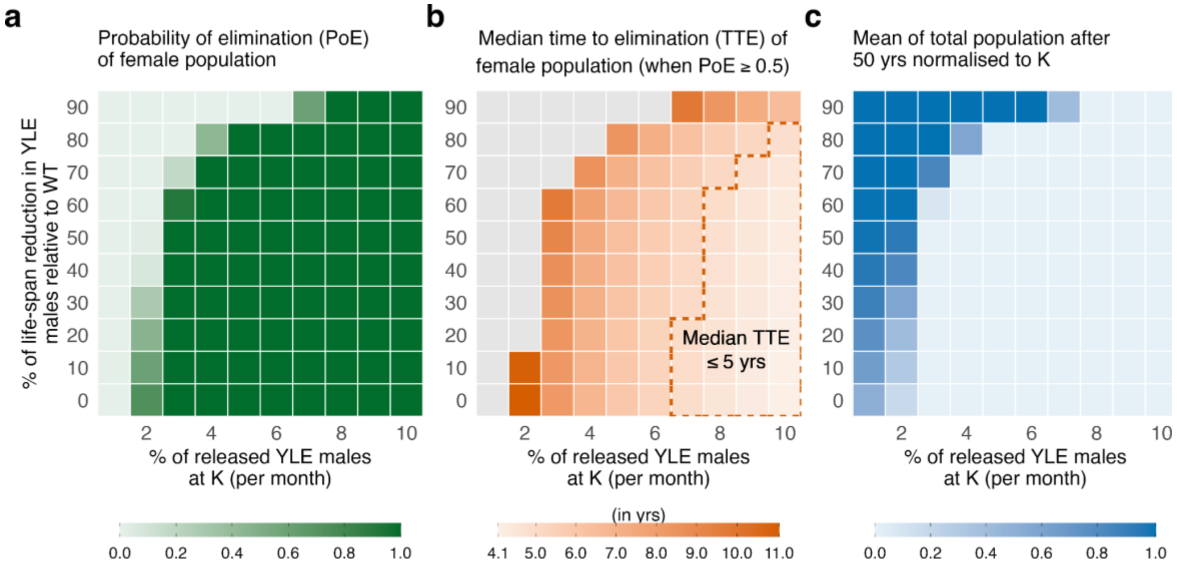


**Fig C. Heatmaps showing the performance of Y-linked editor (YLE) targeting a female-specific dominant lethality gene across release intensity and carrier fitness cost.** Each panel maps outcomes over a common parameter grid where x-axis is the release proportion of YLE males denoted as the percent of adult males at carrying capacity, released monthly for 10 years and the y-axis is the fitness cost to YLE carriers as percent reduction in their lifespan. For each parameter combination, 100 stochastic simulations were performed. Panel **(a)** shows the probability of female elimination (0-1). Panel **(b)** shows the median time to female elimination (in years), computed only when the elimination probability is ≥50%. Darker orange indicates higher time to elimination, and grey color indicates that the elimination probability <50%. The dashed orange perimeter marks parameter combinations where the median time to elimination is ≤5 years. Panel **(c)** shows the final total population after 50 years where population is normalized with respect to the adult carrying capacity, $K$, and averaged across runs (0-1 scale).

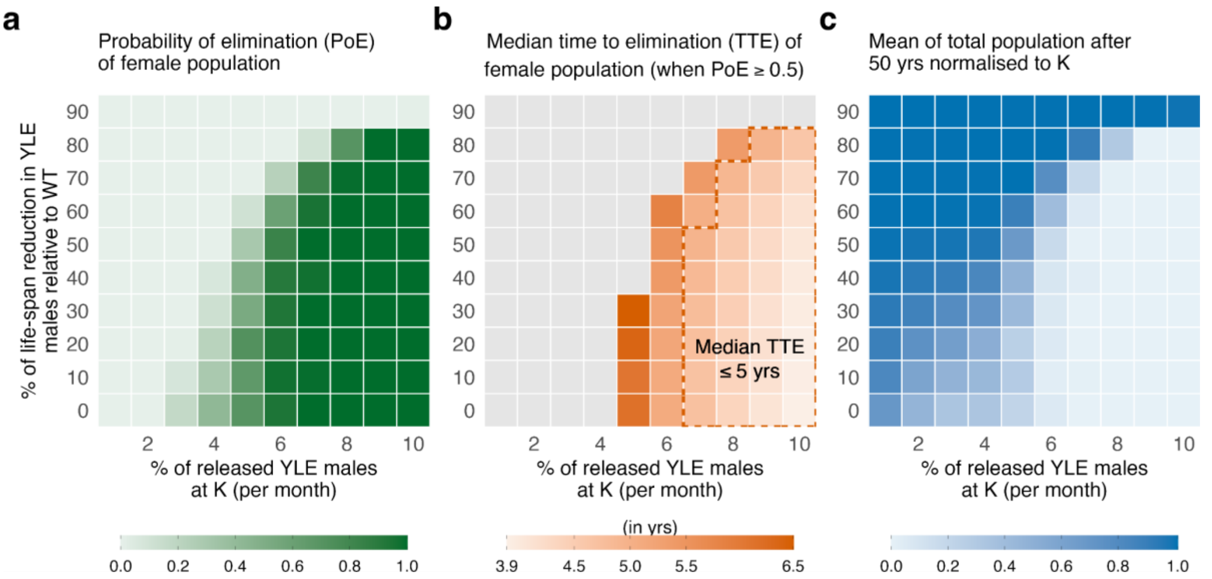


**Fig D. Heatmaps showing the performance of Y-linked editor (YLE) targeting a female-specific dominant lethal gene across release intensity and carrier fitness cost.** Each panel maps outcomes over a common parameter grid where x-axis is the release proportion of YLE males denoted as the percent of adult males at carrying capacity, released monthly for five years and the y-axis is the fitness cost to YLE carriers as percent reduction in their lifespan. For each parameter combination, 100 stochastic runs were performed. Panel **(a)** shows the probability of female elimination (0-100%). Panel **(b)** shows the median time to female elimination (in years), computed only when the elimination probability is ≥50%. Darker orange indicates higher time to elimination, and grey color indicates that the elimination probability <50%. The dashed orange perimeter marks parameter combinations where the median time to elimination is ≤5 years. Panel **(c)** shows the final total population after 50 years where population is normalized with respect to the adult carrying capacity, *K*, and averaged across runs (0-1 scale).

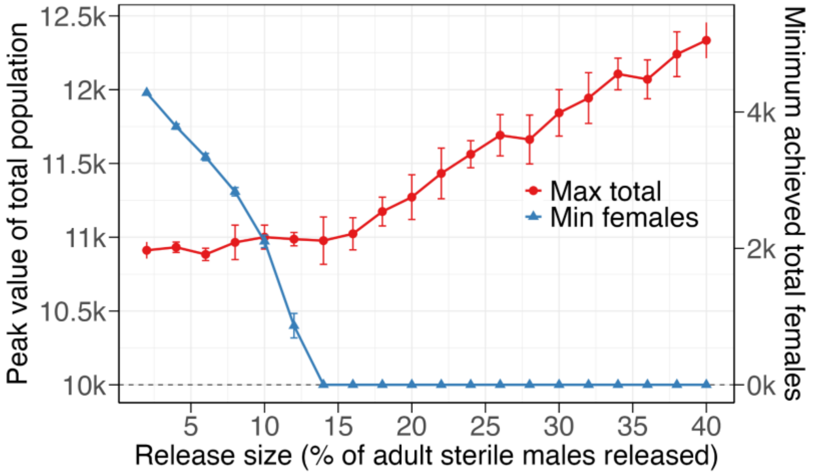


**Fig E. Suppression efficacy and population peaks as a function of release size for release of rodents carrying a female-specific dominant lethal gene (fsRRDL).** Mean of peak total population (red) and minimum wild-type female population (blue) achieved during a ten-year fsRRDL campaign as a function of monthly release size. The dashed dark grey line represents the equilibrium population size in the absence of interventions (*K*=10,000). Data represent means from 100 stochastic runs.

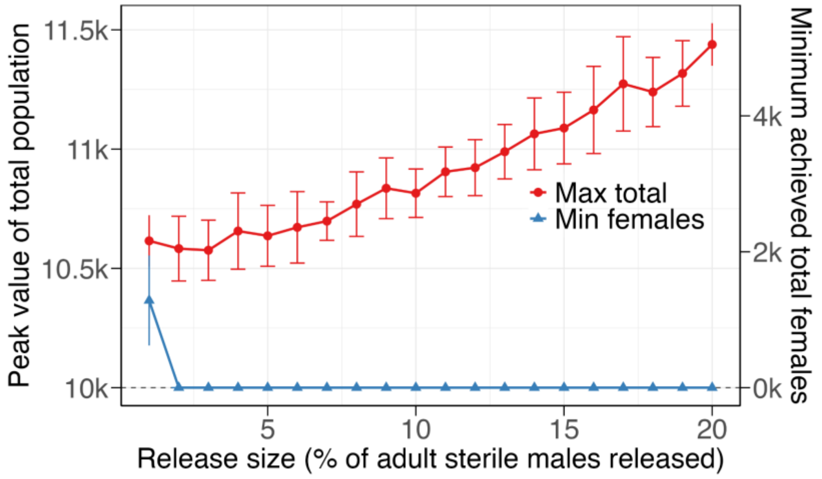


**Fig F. Suppression efficacy and population peaks as a function of release size for a Y-linked editor (YLE).** Mean of peak total population (red) and minimum wild-type female population (blue) achieved during a ten-year YLE campaign as a function of monthly release size. The dashed dark grey line represents the equilibrium population size in the absence of interventions (*K*=10,000). Data represent means from 100 stochastic runs (baseline parameters: **Table B**).

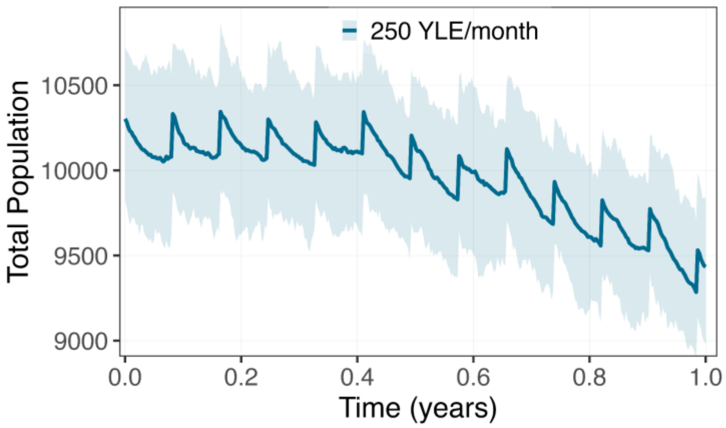


**Fig G. Transient population overshoot during the initial phase of the Y-linked editor (YLE) suppression campaign.** The total mouse population trajectory is shown for the first year of intervention involving continuous monthly releases. A minor, transient increase of approximately 3% above the equilibrium capacity (*K*=10,000) is observed during the first nine releases. This temporary rise persists for ~35 weeks before the population drops below baseline levels and suppression takes effect.

### Supplementary tables

**Table A. Description and baseline values of mouse life history parameters used in the MouseGD framework.** All parameter values are adopted from Brown *et al.* [1].

| Parameter (symbol/name) | Description | Baseline value | Units |
|---|---|---|---|
| $t_{Gest}$ | Gestation duration: average time females remain pregnant before parturition. | 19 | days |
| $t_{Nurs}$ | Nursing duration: time offspring remain dependent on the dam post-birth. | 23 | days |
| $t_{Ado}$ | Adolescence duration: time juveniles remain pre-reproductive before entering the adult stage. | 37 | days |
| $t_{Ad}$ | Mean residence time in the adult stage; determines baseline daily adult mortality. | 690 | days |
| $\mu_{Ad}$ | Density-independent adult mortality applied each day to adults. | 1/690 | per day |
| $L$ | Average number of litters produced per adult female per year; maps to a daily conception probability. | 7.5 | litters/female/yr |
| $Q$ | Mean pups per litter; used as the Poisson mean for offspring at birth events. | 6 | pups per litter |
| $K$ | Adult carrying capacity; regulates the intensity of density-dependent adult mortality. | 10,000 | Adults # |
| $\Theta$ | Shape parameter of density dependence; steeper values increase mortality as adults approach/exceed ($K$). | 22.4 | Unitless |

**Table B. Model parameters for Y-linked editor (YLE) inheritance, fitness, and release strategies.** The table lists baseline values used for primary simulations alongside the parameter ranges used for global sensitivity analysis (SA). U($a$, $b$) denotes a continuous uniform distribution on the interval [$a$, $b$]; parameters marked "fixed" were held constant during sensitivity sweeps. Literature sources for specific parameter estimates are provided in the reference column.

| Para-meter | Description | Baseline value | SA distribution | Reference |
|---|---|---|---|---|
| $\mu$ | Per-allele cleavage rate in (yWW) males during gametogenesis (edit initiation in homozygotes). | 0.97 | U(0.80, 1.00) | [2] |
| $q$ | In (yWW) after cleavage: fraction of repairs that yield resistant or recessive variant (V); remaining (1-q) yield desired edit (E). | 0.00 | U(0.00, 0.20) | [3] |
| $c$ | Cleavage rate of a (W) allele in heterozygous males (yWE/yWV) (edit initiation in heterozygotes). | 0.93 | U(0.80,1.00) | [2] |
| $j$ | Given cleavage in heterozygotes: rate of NHEJ (vs. HDR). With HDR (1-j), the (W) copies the homolog (to E or V). | 0.25 | U(0.00, 1.00) | [4,5] |
| $p$ | In heterozygotes after NHEJ: rate that NHEJ outcome is resistant or recessive lethal (V) (vs. E with prob. (1-p)). | 0.00 | U(0.00, 0.20) | [3] |
| $f_y$ | Relative fitness of YLE males with respect to wild-type males as reduction in expected lifespan | 0.8 | U(0.20, 1.00) | [3] |
| $f_S$ | Expressivity of female sterility with one or more E allele (0=no effect, 1=complete). | 1 (0 when target gene affects viability) | U(0.80, 1.00) | [2] |
| $f_L$ | Expressivity of female lethality with one or more E allele (0=no effect, 1=complete). | 0 (1 when target gene affects viability) | U(0.80, 1.00) | [2] |
| $rel$ | Released proportion of adult males at $K$ introduced as YLE (yEE) males (releases per month). | (scenario-specific) | U(0.01, 0.20) | |

**Table C. Relative influence of model parameters on eradication outcomes.** Sensitivity analysis was performed using boosted regression trees (BRTs) to evaluate the impact of input parameters on two metrics: the probability of successful population eradication and the time to eradication (conditioned on success). Values represent the relative contribution (%) of each parameter to the predictive power of the BRT models. Parameters are sorted in descending order of their influence on the probability of eradication.

| Parameter | Description | Rel. Influence (Probability of Elimination) | Rel. Influence (Time to Elimination) |
|---|---|---|---|
| $rel$ | Release Size | 86.99% | 91.98% |
| $f_y$ | YLE Fitness | 11.95% | 1.38% |
| $p$ | Het. Resistance | 0.56% | 3.06% |
| $c$ | Het. Cleavage | 0.18% | 0.57% |
| $q$ | Hom. Resistance | 0.11% | 0.40% |
| $\mu$ | Hom. Cleavage | 0.09% | 0.09% |
| $f_S$ | Fem. Sterility | 0.08% | 1.92% |
| $j$ | NHEJ Fraction | 0.04% | 0.60% |

**Table D. Drive mechanics and release parameters for homing gene drives (HGDs).** Parameter sets are distinguished for "ideal" scenarios (assuming perfect homing and no resistance) and "non-ideal" scenarios (incorporating observed rates of non-homologous end-joining (NHEJ) repair and formation of functional resistance alleles). Sources for parameter values are listed in the references column.

| Parameter | Description | Ideal condition | Non-ideal condition | Reference |
|---|---|---|---|---|
| p_C | Rate that the target allele is cut in a germline cell (cutting efficiency) | 1.00 | 0.97 | [2] |
| p_N | Rate of NHEJ repair given that a cut occurs | 0.00 | 0.25 | [4,5] |
| p_L | Fraction of NHEJ events that generate a loss-of-function (disruptive) allele | 1.00 | 5/6 | [6,7] |
| f_W | Relative fitness of wild-type allele (W) | 1.00 | 1.00 | [3] |
| f_H | Relative fitness of homing drive allele (H) | 1.00 | 0.80 | [3] |
| f_R | Relative fitness of functional resistant allele (R) | 1.00 | 1.00 | [3] |
| f_N | Relative fitness of non-functional (loss-of-function) allele (N) carriers with respect to wild-type as reduction in expected lifespan | 0.00 | 0.50 | |
| rel | Released adult males (yHH) males (released once) | 250 | 250 | |

**Table E. Drive mechanics and release parameters for Y-linked X-shredder systems.** Values define "ideal" performance (perfect sex-ratio distortion) versus "non-ideal" performance (accounting for incomplete shredding efficiency and associated fitness costs). Relevant literature sources for these values are cited in the references column.

| Paramete r | Description | Ideal condition | Non-ideal condition | Reference |
|---|---|---|---|---|
| c_X | Probability that an X chromosome is successfully shredded in X-shredder males | 1.00 | 0.97 | [2] |
| c_rX | Rate of resistance chromosome generation | 0.00 | 0.25 | [4,5] |
| f_XA | Relative fertility fitness of male carrying attacking Y chromosome or X shredder construct (mXA, mRA) | 1.00 | 0.80 | [3] |
| rel | Released adult males (mXA) with X shredder construct (released once) | 250 | 250 | |
